# Food recognition in a blood-feeding insect: characterization of the pharyngeal taste organ

**DOI:** 10.1101/2022.11.15.516640

**Authors:** Isabel Ortega-Insaurralde, José Manuel Latorre-Estivalis, Andre Luis Costa-da-Silva, Agustina Cano, Teresita C. Insausti, Hector Salas Morales, Gina Pontes, Martín Berón de Astrada, Sheila Ons, Matthew DeGennaro, Romina B. Barrozo

## Abstract

**Background:** Obligate blood-feeding insects obtain the nutrients and water necessary to ensure survival from the vertebrate blood. The internal taste sensilla, situated in the pharynx, evaluate the suitability of the ingested food. Here, through multiple approaches, we characterized the pharyngeal organ (PO) of the hematophagous kissing bug *Rhodnius prolixus* to determine its role in food assessment. The PO, located antero-dorsally in the pharynx, comprises 8 taste sensilla that become bathed with the incoming blood.

**Results:** We showed that these taste sensilla house gustatory receptor neurons projecting their axons through the labral nerves to reach the subesophageal zone in the brain. We found that these neurons are electrically activated by relevant appetitive and aversive gustatory stimuli such as NaCl, ATP and caffeine. Using RNA-Seq, we examined the expression of sensory-related gene families in the PO. We identified gustatory receptors, ionotropic receptors, transient receptor potential channels, pickpocket channels, opsins, takeouts, neuropeptide precursors, neuropeptide receptors and biogenic amine receptors. RNA interference assays demonstrated that the pickpocket channel *Rproppk014276* is necessary for salt detection during feeding.

**Conclusion:** We provide evidence of the role of the pharyngeal organ in food evaluation. This work shows the first comprehensive characterization of a pharyngeal taste organ in a hematophagous insect.

## Background

Contact chemoreception or taste is the sensory modality that allows an animal to detect non-volatile chemicals on a substrate or in a liquid medium. Among other functions, such as identifying conspecifics and/or oviposition sites, the taste system allows animals to predict the nutritional or toxic quality of the food even before feeding on it. Thus, the taste sense drives the final feeding decisions of animals that will undoubtedly have determinant physiological consequences in their lives (Muñoz et al., 2020).

Food evaluation in insects is a highly relevant task performed by specialized peripheral sensory organs and their functional units, the taste sensilla. At this stage, the presence or absence of chemical components of the food leads to the final decision: to eat or not to eat. Gustatory receptor neurons (GRNs) represent the cellular platform of the taste system housed within the taste sensilla (Liman et al., 2014). Embedded in the membranes of GRNs, sensory protein receptors are responsible for the molecular detection of taste stimuli. Activation of these GRNs evokes an electrical signal that travels to the insect brain where the decision to accept or reject the food is triggered.

A sequence of feeding events starts once a blood-feeding insect reaches a vertebrate host, as has been described for the kissing bug *Rhodnius prolixus* (Barrozo, 2019). First, the blood feeder gustatory sense assesses the skin and if no aversive stimuli are detected, the insect pierces the skin (Pontes et al., 2014; 2022). Second, before starting the true ingestion, pharyngeal and cibarial muscles located in the head produce contractions, sucking a small quantity of blood. The insect probes the incoming food by evaluating its quality (Pontes et al., 2017). If the ingested blood fulfills the insect’s feeding requirements, the animal continues the feeding behavior; if not, the animal leaves the host and searches for another (Barrozo, 2019; Smith and Friend, 1970).

Gustatory blood assessment seems to occur in internal taste sensilla located strategically along the pharynx, such as the papillae on the cibarium of *Aedes aegypti* mosquitoes and species of the genera *Culex sp., Culiseta sp*. and *Anopheles sp*., the short-peg sensilla of the pharyngeal organ (PO) of the kissing bugs *R. prolixus* and *Triatoma infestans* and the labro-cibarial sensilla of *Simulium spp*. (Barth, 1952; Bernard, 1974; Jefferies, 1987; Lee and Craig, 1983; Owen, 1963; Pontes et al., 2014; Williams and Savage, 2009). All these sensilla are bathed with blood as soon as the first sip passes through the alimentary canal.

Few chemicals present in the ingested blood have been shown to induce blood-feeding in insects, and the identification of these components could lead to the development of anti-feedants (Barrozo, 2019). Adenosine nucleotides are the main known phagostimulant compounds in the vertebrate blood. They are released by lysis of erythrocytes, platelets and epithelial cells as a damage response to the piercing or severing during blood intake (Born and Kratzer, 1984; Friend and Smith, 1977; Forsyth et al., 2011). Several insects are responsive to ATP, including kissing bugs, some mosquitoes, tsetse flies and fleas, while others respond better to ADP (Barrozo, 2019). Low concentrations of salts, necessary to maintain the endogenous salt balance, have also been proved to elicit appetitive behaviors in kissing bugs, mosquitoes, fleas and bed bugs (Galun, 1966; Jové et al., 2020; Liscia et al., 1993; Pontes et al., 2017; Romero and Schal, 2013). On the other hand, molecules perceived as bitter by humans like quinine, caffeine and quinidine have an antifeedant effect in mosquitoes such as *An. gambiae* and *Ae. aegypti* (Dennis et al., 2019; Ignell et al., 2010; Kessler et al., 2013, 2014). Furthermore, the detection of quinine, caffeine, berberine, salicin or high NaCl (> 0.2 M) inhibits feeding in *R. prolixus* (Asparch et al., 2016; Cano et al., 2017; Pontes et al., 2014, 2017).

*Rhodnius prolixus* is a competent vector of the Chagas disease that transmits the protozoan parasite *Trypanosoma cruzi* to humans through its feces following a blood meal. This neglected parasitic disease affects several countries in Latin America (WHO, 2020). Under this context, taste perception represents a relevant target for suppressing blood feeding and, subsequently the pathogen transmission. Morphological studies have shown taste sensilla on the anterodorsal region of the pharynx of kissing bugs. These sensilla have been postulated as food sensors (Barth, 1952; Bernard, 1974; Pontes et al., 2014). However, the sensory basis of blood gustatory assessment by internal taste organs, such as the PO, has been barely studied in blood-feeding insects (Barrozo, 2019; Benton, 2017). We provide neuroanatomical, physiological, and molecular evidence uncovering the role of the PO in food assessment in *R. prolixus* through a multi-approach analysis. We found that taste sensilla in the PO house gustatory receptor neurons (PO-GRNs), which respond to gustatory stimuli relevant to kissing bugs, such as NaCl, ATP, and caffeine. The taste information then reaches the subesophageal zone in the brain via the labral nerves, where the information is likely first processed by the CNS as in other insects (Harris et al., 2015; Ignell and Hansson, 2005). Furthermore, a vast repertoire of genes is expressed in the PO, in which several candidates may be related to food recognition functions. Among them, the pickpocket channel receptor *Rproppk014276* is shown to be involved in salt detection, salts being crucial stimuli to trigger feeding in *R. prolixus*.

## Results

### The Pharyngeal Organ (PO) houses GRNs

The PO consists of 8 short-peg sensilla located along 80 μm in the anterodorsal region of the pharynx of *R. prolixus* as shown in the scheme of **Fig. 1A**. Dispersedly distributed, these uniporous sensilla are candidates to sense the incoming blood meal **(Fig. 1A)**. Histological transverse sections of the pharynx allowed the recognition of winding dendrites inside the sensilla **(Fig. 1B, C)**. Figure 1C shows a sensory neuron breaking through the thick cuticle of the dorsal wall of the pharynx, a candidate for being a PO gustatory receptor neuron responsible for food sensing.

**Figure 1.**
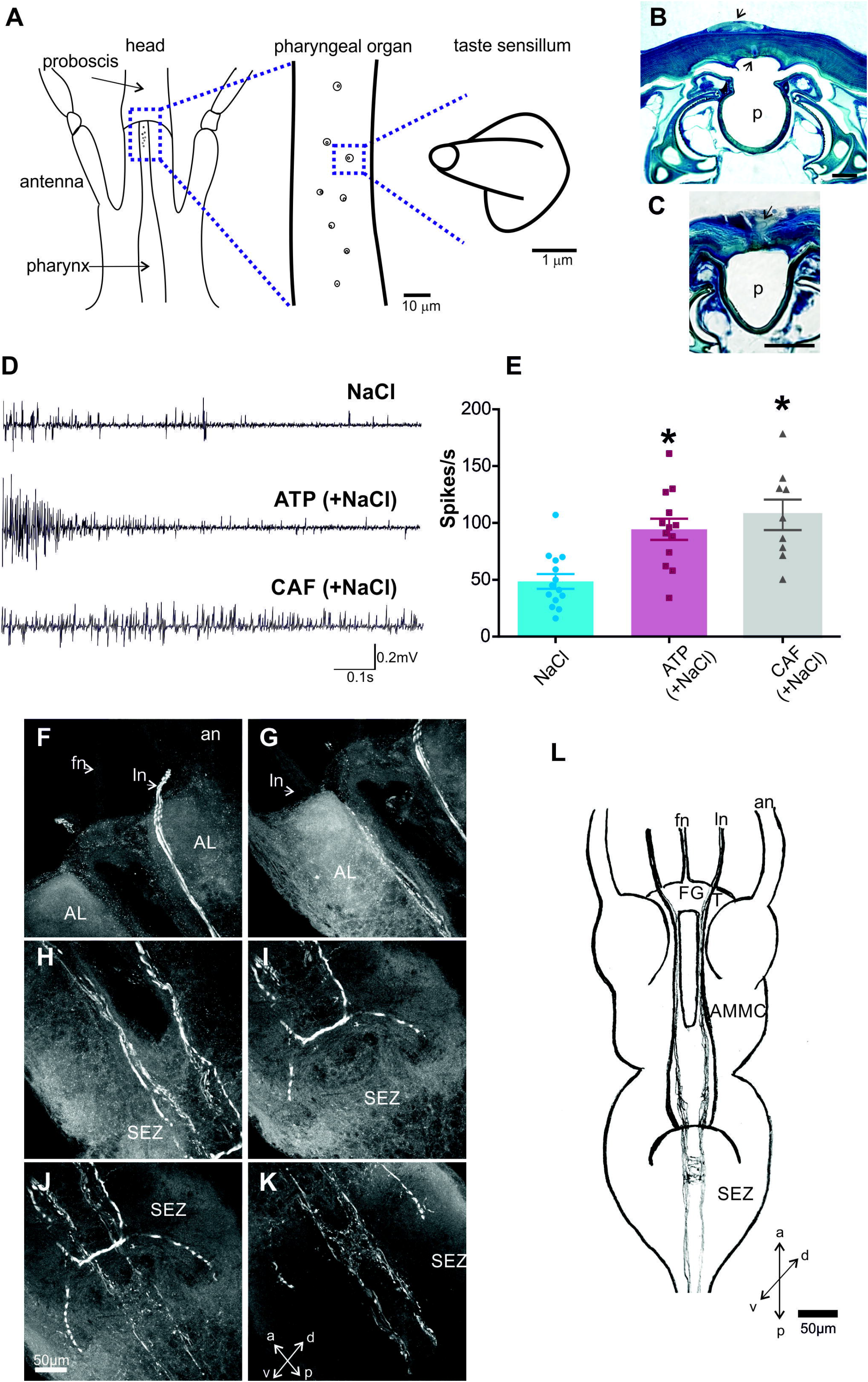
Morphology and function of taste sensilla of the PO. **(A)** Schemes of the PO and the single uniporous sensilla at different scales. A longitudinal and ventral view of the head and pharynx, unveiling the 8 taste sensilla of the PO. The PO is located anterodorsally to the head and pharynx. **(B-C)** Photographs of histological cross sections of the PO. **(B)** Dorsally to the pharynx (p), indicated with arrows, a taste sensillum and 2 cell bodies can be identified. Horizontal bar denotes 2 μm **(C)** A dendrite of a putative PO-GRN breaking through the pharyngeal cuticle is observed, shown with an arrow. Horizontal bar indicates 10 μm. **(D-E)** Electrophysiological responses of PO-GRNs. **(D)** Example of spike discharges of the GRNs housed in the PO. Typical responses to NaCl, ATP and caffeine are shown. **(E)** Scatter plots are shown and bars represent the mean firing responses (mean ± s.e.m.) of PO-GRNs to the 3 gustatory stimuli. ATP and caffeine produced significantly more action potential discharges than NaCl. Asterisks indicate statistical differences between insects stimulated with ATP or caffeine versus those stimulated with NaCl alone (p < 0.0005). ATP and caffeine solutions also contained NaCl (see Methods). 9 to 14 replicates were carried out *per* treatment. **(F-L)** Anterograde staining of the PO-GRNs. **(F-K)** Photographs at different levels of the brain of insects, showing 2 subtypes of PO-GRNs reaching the subesophageal zone (SEZ). One subtype constituted by thick axons showed to project contralaterally in the SEZ exclusively. Another subtype of PO-GRNs arborise ipsilaterally at the medial and dorsal region of the SEZ, and continue to the prothoracic ganglion. **(L)** Reconstruction of the arborizations of PO-GRNs that reach the brain. fn: frontal nerve, ln: labral nerve, an: antennal nerve, FG: frontal ganglion, T: tritocerebrum, AL: antennal lobe, AMMC: antennal motor mechanosensory center, SEZ: subesophageal zone. a: anterior, p: posterior, d: dorsal, v: ventral.

Key taste components of the ingested food might be detected in the PO-GRNs. Therefore, the functional role of PO-GRNs in food detection was confirmed through electrophysiological recordings **(Fig. 1D, E)**. The electrophysiological activity of these sensory neurons was recorded by placing the recording electrode next to the PO. We recorded the global response of the PO-GRNs upon gustatory stimulation with NaCl, ATP and caffeine **(Fig. 1D, E)**, known the three to produce appetitive or aversive feeding responses in *R. prolixus*. In all the recordings, we found a neuronal activity for the 3 stimuli **(Fig. 1D)**. Even though several responding neurons became activated, spike sorting of the responses of individual neurons was difficult. Nevertheless, general responses were quantified upon insect stimulation with the different tastants **(Fig. 1E)**. Statistical differences were found among the gustatory stimuli (K-W = 15.3, p = 0.0005, *post hoc* Dunn’s comparisons against NaCl group p < 0.05). NaCl elicited action potentials (48.6 ± 6.5) upon stimulation, whereas the overall firing activity was significantly increased with ATP (94.5 ± 9.3) or caffeine (107.2 ± 13.4). These results showed that the sensilla present on the PO effectively house sensory neurons sensitive to behaviorally relevant stimuli, demonstrating a role for the PO during taste assessment of incoming food in *R. prolixus*.

### The PO-GRNs project into the subesophageal zone

Following stimulus detection, taste information must reach the brain for integration and processing. Therefore, and through anterograde backfills of PO-GRNs, we examined the morphology, topography and sites of arborizations of these sensory neurons in the central nervous system. Five preparations showed stained neuronal arborizations in the brain, which showed similar results. Two morphotypes or subtypes of GRNs were recognized. Thus, the axons of these neuronal subtypes extend from the PO via the labral nerves **(Fig. 1F-K)**. A bundle of thin subtype of PO-GRNs **(Fig. 1F-K)** arborize first ipsilaterally at the medial and dorsal region of the subesophageal zone (SEZ), and continue to the prothoracic ganglion **(Fig. 1J-K)**. Another subtype of PO-GRN was easily recognized due to its thickness regarding the other subtype **(Fig. 1I-J)**. Interestingly, these subtype of neurons showed to project contralaterally in the SEZ exclusively, as indicated in the photo **(Fig. 1I-J)** and in the reconstruction **(Fig. 1L)**. These results showed the SEZ as the first central relay of PO-GRNs in *R. prolixus*.

### The PO expresses several sensory-related gene families

RNA sequencing (RNA-Seq) allowed us to examine the gene expression of the isolated PO. We found representatives of gene families associated with sensory functions previously characterized in other insects **(Fig. 2)**. Several of them possess putative gustatory functions relevant at the time of food recognition **(Fig. 2, 3)**. A total of 307,059,020 raw reads were produced from three libraries. After quality control and trimming, a total of 91.1M, 75.7M and 78.9 M of cleaned reads were obtained for the three replicates, respectively. A total of 53.5M, 42.4M and 49.1M reads mapped to the *R. prolixus* genome. Gene expression data of the main sensory-related gene families represented as Log_10_ (TPM (Transcripts *Per* kilobase *per* Million read) +1), were obtained **(Fig. 2, 3)**. Among others, we found sensory genes representative of seven families that include: gustatory receptors (GRs), ionotropic receptors (IRs), pickpocket channels (PPKs), transient receptor potential channels (TRPs), opsins (Ops), odorant-binding proteins (OBPs) and chemosensory proteins (CSPs) **(Fig. 2)**. Takeouts (TOs), neuropeptide precursors (NPs), neuropeptide receptors (NPRs) and receptors of biogenic amines (BARs) were also detected, many of them were known to take part in the regulatory mechanisms of the feeding behavior in several insect models **(Fig. 3)** (Sarov-Blat et al., 2000; Dacks et al., 2003; Yarmolinsky et al., 2009; Sterkel et al., 2011).

**Figure 2.**
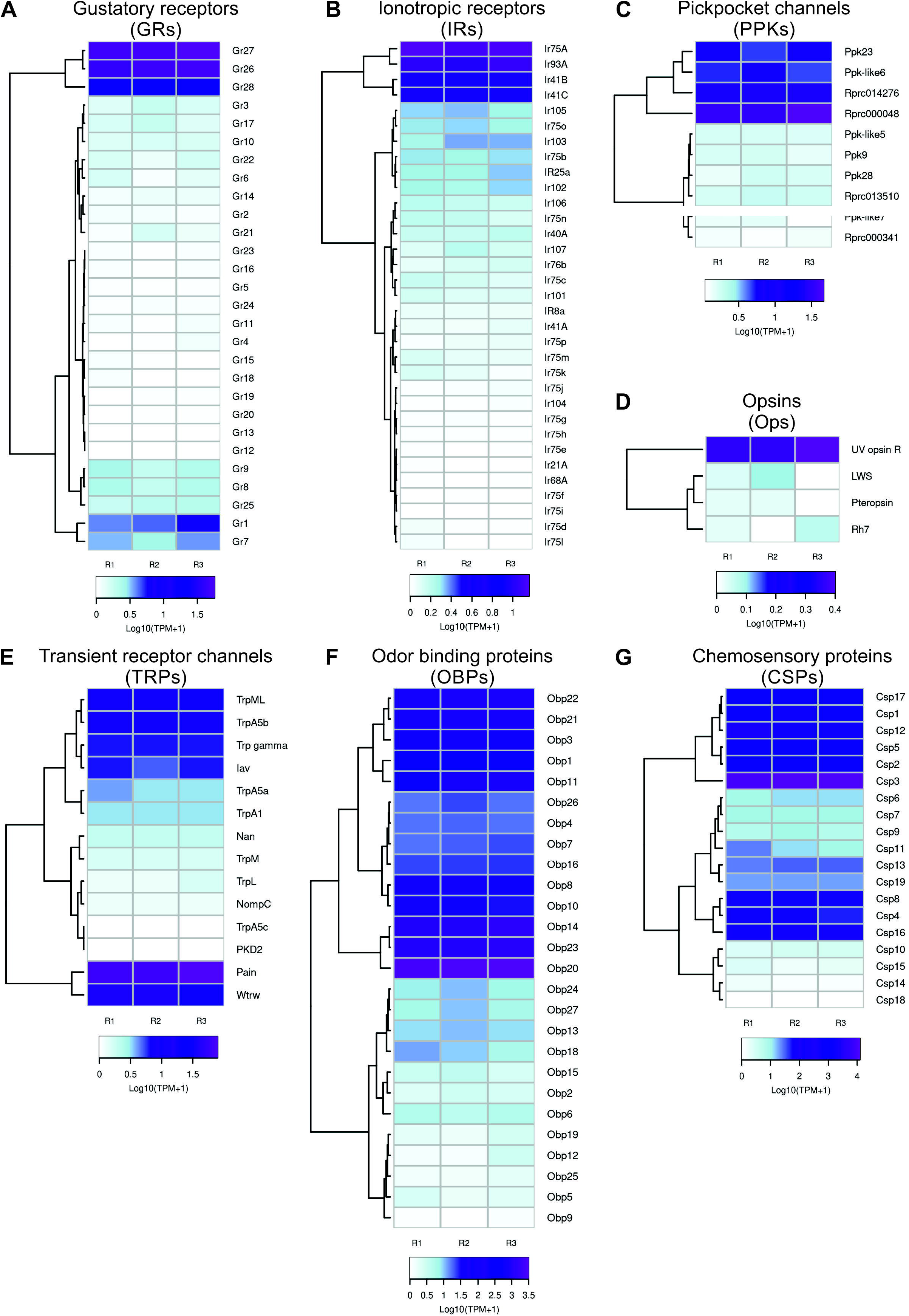
Expression of sensory-related genes in the PO. Heatmaps were built using Log_10_ (Transcripts *per* kilobase *per* Million read - TPM +1) as input of the gplot R package. Transcript abundance was represented in a color scale where white/purple represents the lowest/highest expression. A dendrogram was plotted using hierarchical clustering of gene expression values based on Euclidean Distances and a complete linkage method for clustering. **(A)** gustatory receptors (GRs) **(B)** ionotropic receptors (IRs) **(C)** pickpocket ion receptor channels (PPKs) **(D)** opsins (Ops) **(E)** transient receptor potential channels (TRPs) **(F)** odorant-binding proteins (OBPs) **(G)** chemosensory proteins (CSPs)

Overall, a low expression of GRs can be noticed **(Fig. 2A)**. This was an unexpected result since GRs are mostly associated with a primary and conserved role in the taste sense of insects (Montell, 2009). However, a few GR encoding genes showed enhanced expression, such as the *RproGr26, RproGr27* and *RproGr28*. Besides, *RproGr1*, the orthologous gene of the fructose receptor *DmelGr43a* (Freeman et al., 2014), also showed relatively high expression level.

Regarding IR expression, high levels for *RproIr93a* and *RproIr75a* were detected **(Fig. 2B)**. Yet, and to a lesser extent, high expression was observed for *RproIr41b* and *RproIr41c* **(Fig. 2B)**. IRs are ligand-gated ion channels derived from variant ionotropic glutamate receptors that can function as chemo-, hygro- and thermoreceptors (Croset et al., 2010; Rytz et al., 2013). It is feasible that these candidates possess similar roles in *R. prolixus*.

PPKs are a family of amiloride-sensitive degenerin/epithelial sodium channels (DEG/ ENaC) that mediate salt and water sensation, among other functions (Kellenberger and Schild, 2002; Zelle et al., 2013). Interestingly, from the 10 PPK-encoding genes present in the genome of *R. prolixus, RPRC000048, Rproppk014276, Rproppk23* and *Rproppk-like6*, exhibited high expression **(Fig. 2C)**. This is similar to what occurs in the antennal sensilla for the *Rproppk014276* (Pontes et al., 2022), this PPK could be an interesting candidate to play a role in salt detection within the PO.

Among the 4 opsins that were found to be detected in the PO, *RproUVopsin* was the most expressed **(Fig. 2D)**. Opsins are known to be involved in photo-, mechano- and thermosensation (Leung and Montell, 2017). More recently, these receptors were demonstrated to behold a gustatory function in the *D. melanogaster* labellum (Leung et al., 2020). *RproRh7*, an orthologous of *DmelRh7* in fruit flies, was related to the detection of the aversive compound aristolochic acid (Leung et al., 2020). This is the first report of opsins expression in the gustatory tissue of a blood-sucking insect.

TRPs are a superfamily of cation-conducting membrane proteins involved in mechano-, chemo-, photo- and thermosensation (Damann et al., 2008; Venkatachalam and Montell, 2014). Regarding the TRP repertoire expressed in the PO tissue, an enhanced expression of *RproPainless* and *RproWaterwitch* followed by *RproTrpml, RproTrpa5b, RproTrp-gamma* and *RproInactive* were observed **(Fig. 2E)**. This highly conserved sensory gene family might represent putative sensors of thermal, nociceptive and bitter cues in the alimentary canal.

High expression levels of OBPs-encoding genes in the PO were also identified. It is noteworthy the high expression levels of *RproObp20, RproObp14* and *RproObp23* **(Fig. 2F)**. Moreover, we found high expression of CSP-encoding genes in the PO, especially *RproCsp3, RproCsp5, RproCsp2, RproCsp1, RproCsp17*, and *RproCsp12* **(Fig. 2G)**. OBPs and CSPs are soluble proteins that bind small molecules, such as odorants and pheromones, in the lymph of insect chemosensilla (Pelosi et al., 2018) among other functions in non-sensory tissues (Rihani et al., 2021). The relative enhanced expression of OBPs and CSPs in the PO heavily indicates them as attractive candidates to aid in the mechanisms of food evaluation.

The TOs are circadian regulated genes whose expressions are induced by starvation and act as hormone carriers (Meunier et al., 2007, Sarov-Blat et al. 2000, Du et al. 2003). TOs are thought to regulate the peripheral sensitivity to phagostimulant stimuli under starvation conditions and male courtship behavior (Bohbot and Vogt, 2005; Fujikawa et al., 2006; Sarov-Blat et al., 2000; Meunier et al., 2007; Dawaulder et al., 2002). They are usually expressed in organs such as labella, crop, antenna and brain. We found that TOs encoding-transcripts are also detected in the PO. *RproTo2* and *RproTo1* showed the highest expression concerning other members of the same family (*RproTo4, RproTo6, RproTo7, RproTo14* and *RproTo15*) which also presented high levels of relative expression **(Fig. 3A)**. High expression levels of biogenic amine receptors (BARs) were also detected, particularly the 5-hydroxytryptamine 2B receptor (*Rpro5HTR2b*), the acetylcholine receptor C (*RproAChRc*) and the dopamine/ecdysone receptor (*RproDopEcR*) **(Fig. 3B)**. The *Rpro5HTR2b* is activated by 5-hydroxytryptamine, i.e. serotonin; and *R. prolixus* depends on this serotonergic system to successfully complete the feeding behavior (Paluzzi et al., 2015). Interestingly, this is the first report of the presence of the *AChRc* in a gustatory organ of any organism studied so far. *DopEcR* is activated by dopamine and ecdysteroids, such as ecdysone and 20-hydroxyecdysone (Srivastava et al., 2005; Kang et al., 2019). Serotonin, dopamine and 20-hydroxyecdysone are known to control and regulate feeding in insects (Inagaki et al., 2012; Vleugels et al., 2015; Selcho and Pauls, 2019; Kang et al., 2019).

**Figure 3.**
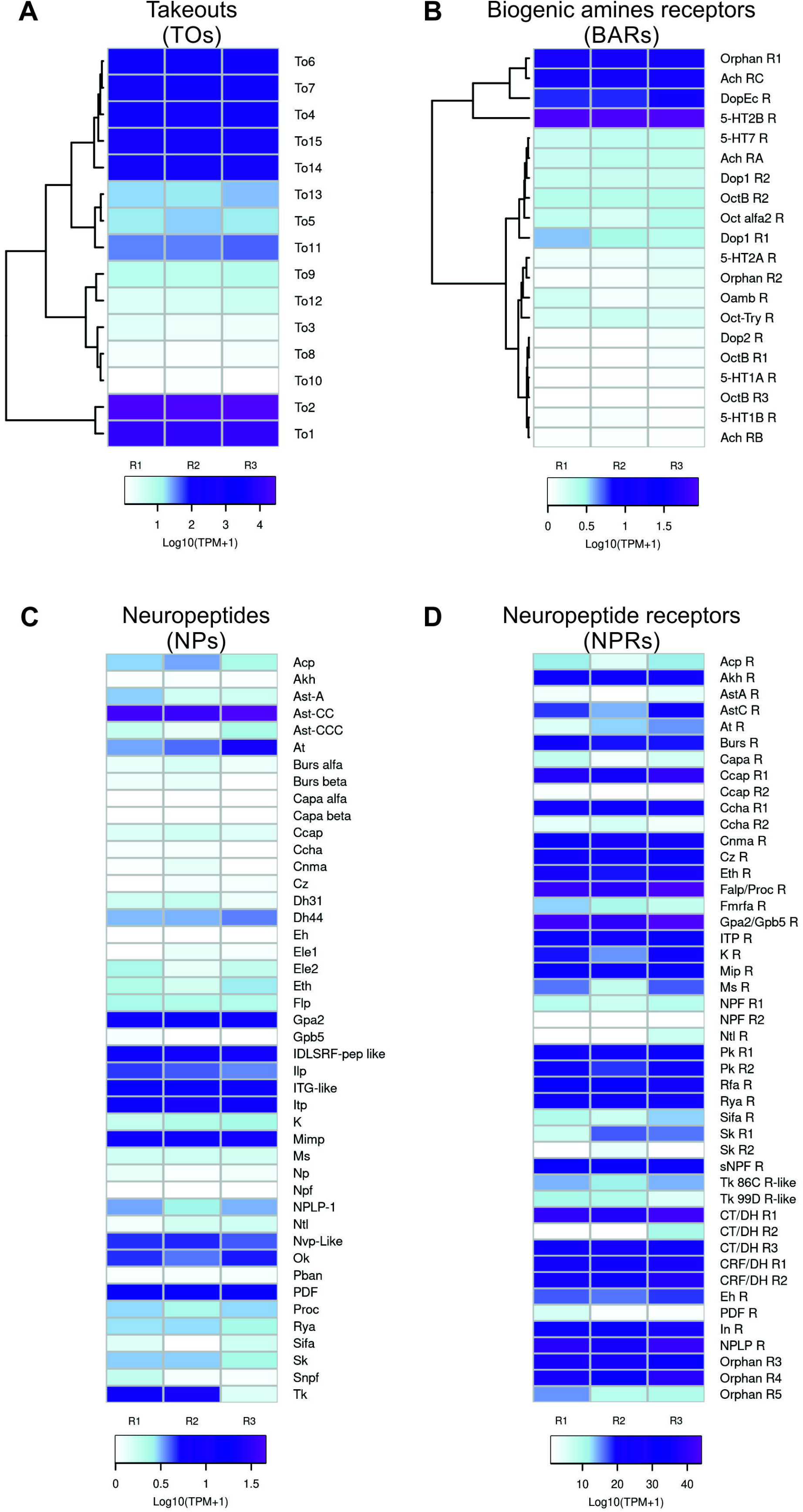
Expression of neuromodulatory genes in the PO. Heatmaps were created using Log_10_ (Transcripts *per* kilobase *per* Million read-TPM +1) as input of the gplot R package. Transcript abundance was represented in a color scale where white/purple represents the lowest/highest expression. A dendrogram was plotted using hierarchical clustering of gene expression values based on Euclidean Distances and a complete linkage method for clustering. **(A)** takeout genes (TOs) **(B)** biogenic amine receptors (BARs) **(C)** neuropeptides (NPs) **(D)** neuropeptides receptors (NPRs)

Transcripts encoding neuropeptide precursors (NPs) and neuropeptide receptors (NPRs) were also identified in the PO, however, their expression can range from low or moderate to high levels **(Fig. 3C, D)**. High expression of both the precursor and receptor was detected only for the heterodimeric glycoprotein hormone *RproGPA2/GPB5*, suggesting a paracrine regulation in the PO **(Fig. 3C)**. High expression of the NP precursor genes allatostatin-CC (*RproAst-CC), RproITG-like*, ion transporter peptide (*RproITP*), myoinhibiting peptide (*RproMIP), RproDLSRF-like* peptide and pigment-dispersing factor (*RproPDF*) was found, even though the expression of their receptors is low, pointing to a hormonal release from the PO to act in an/other tissue(s) **(Fig. 3D)**. Conversely, the proctolin receptor (*RproFalp/ProcR*), the calcitonin-like/ diuretic hormone receptor 1 (*RproCT/DHR1*), the neuropeptide-like receptor 1 (*RproNPLPR*), the crustacean cardioactive peptide receptor 1 (*RproCCAPR1*), the CRF-like/diuretic hormone receptor 2 (*RproCRF/DhR2*), and the insulin receptor (*RproInR*) are highly expressed in the PO. However, the expression of their precursors encoding their peptide ligands are low **(Fig. 3D)**. This could indicate endocrine responses in the PO to neuropeptides released to the hemolymph from other insect tissues. Several of these NPs and NPRs were previously demonstrated to regulate feeding-related events (Nässel and Zandawala, 2019; Ons, 2017).

### Disruption of Rproppk014276 prevents feeding

We decided to investigate the role of the PO in food assessment by disrupting the expression of candidate genes expressed in PO. We focused on *Rproppk014276* and *Rproppk28* because of their known roles in salt detection in *R. prolixus* (Pontes et al., 2022). Knowing that NaCl, at an equivalent amount of the vertebrate plasma, is one of the main phagostimulants of *R. prolixus* and, consequently, crucial to trigger feeding (Cano et al., 2017; Pontes et al., 2017), we predicted that the knock-down of one or both PPKs could prevent this response.

RNA interference experiments to silence *Rproppk014276* (VectorBase RPRC014276) and *Rproppk28* (VectorBase RPRC000471) expression were carried out. Therefore, the four experimental groups consisted in: uninjected insects, dsRNA-ctrl (control group of dsRNA injection), dsRNA-Rproppk014276 and dsRNA-Rproppk28 injected insects. The RT-qPCR results showed effective knock-down of *Rproppk014276* and *Rproppk28* transcript levels in the antennae of *R. prolixus* (**Fig. S1**). Then, the feeding response of control and knocked-down insects was tested to an appetitive solution composed of NaCl and ATP. Note that, uninjected kissing bugs need both NaCl and ATP to elicit a full appetitive response (Pontes et al., 2017). By using an artificial feeder coupled to an electromyographic recording system (**Fig. 4A**), we recorded the activity of the muscles associated with feeding in all experimental groups. Additionally, a group uninjected insects were offered with a feeding solution containing only ATP but not NaCl (no-salt in **Fig. 4**). The rationale behind this assay was to show that kissing bugs do not feed on a salt-free solution, even if ATP is present. Therefore, if *Rproppk014276* and/or *Rproppk28* are involved in salt detection, we would expect that its after knockdown, bugs behave like uninjected insects offered with a salt-free solution.

**Figure 4.**
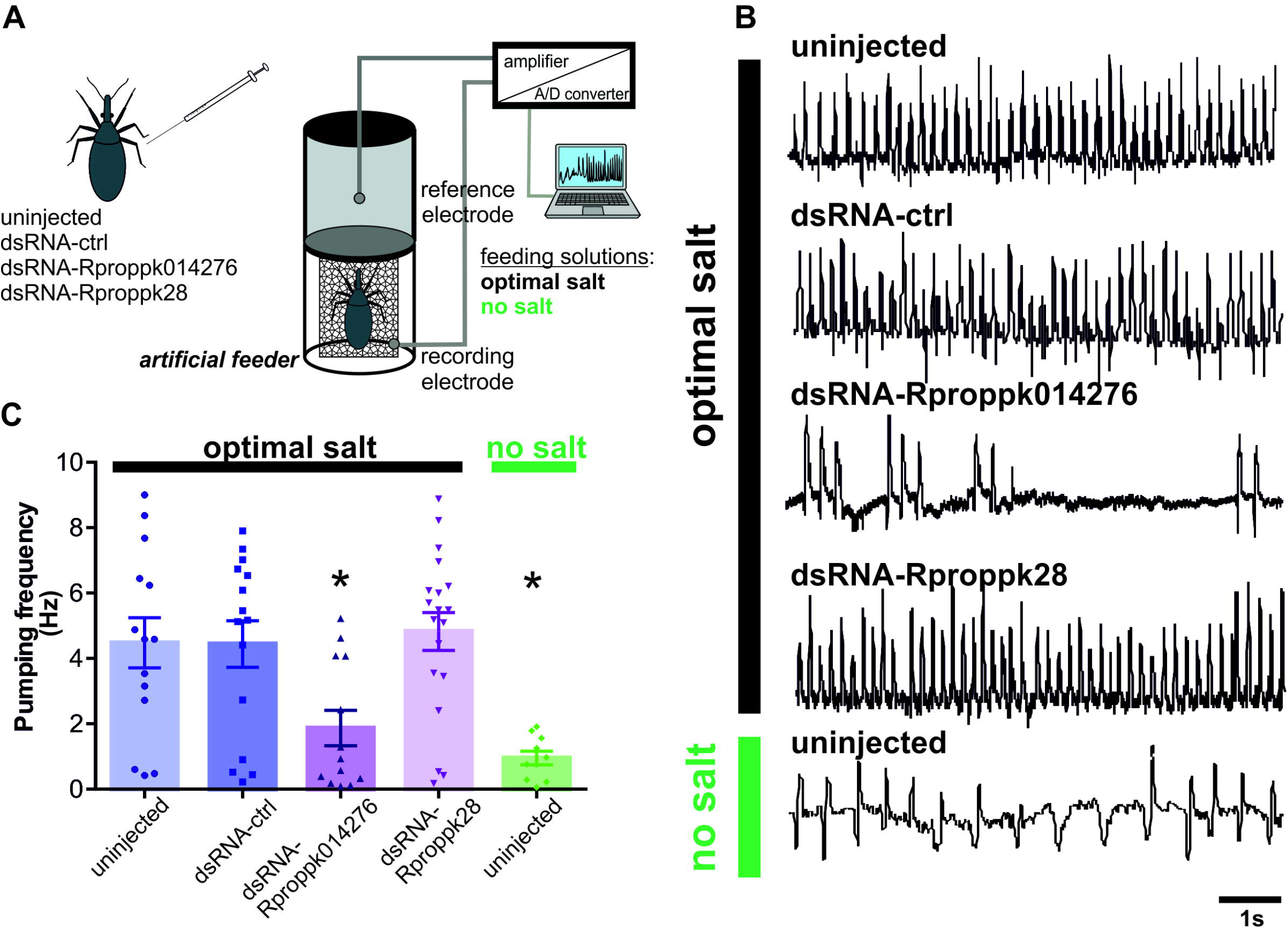
Role of salt-related PPKs during feeding. **(A)** Experimental design for feeding experiments. The feeding activity of 4 groups of insects were tested in the artificial feeder: uninjected, dsRNA-ctrl, dsRNA-Rproppk014276 and dsRNA-Rproppk28. The artificial feeder was coupled to an electromyographic recording system, in which the recording electrode was connected to a metallic mesh where the insect was posed, and the reference electrode was immersed in the feeding solution. Once the insect inserted its mouthparts into the feeder, the electrical circuit was closed generating a base conductance. Changes in the baseline signal were attributed to contractions produced by the sucking muscles when the insect began to feed. Recorded signals were amplified, digitized and stored in a PC. The 4 experimental groups were offered with an appetitive optimal-salt solution. A control group of uninjected insects was tested to a non-appetitive salt-free solution (no salt). **(B)** Example of electromyographic recordings of the muscles involved in feeding. Uninjected, dsRNA-ctrl and dsRNA-Rproppk28 groups showed similar responses to the optimal salt-solution, showing a regular pumping behavior. dsRNA-Rproppk014276 insects offered with optimal salt or uninjected insects offered with no-salt solution exhibited low and scattered pumping activities relative to the other groups. **(C)** Pumping frequency during feeding of treated and uninjected insects on the optimal salt or on the salt-free (no salt) solutions. No differences were found between uninjected, dsRNA-ctrl and dsRNA-Rproppk28 groups. dsRNA-Rproppk014276 insects, instead, showed significantly lower frequency values than the other groups. Note that the dsRNA-Rproppk014276 group offered with optimal salt showed no differences from uninjected insects fed on a salt-free solution. Scatter plots are shown and bars represent the mean pumping frequency (mean ± s.e.m). Asterisks indicate significant differences of each group to uninjected insects fed on the optimal-salt solution (Kruskal-Wallis test, Dunn’s *post hoc* comparisons, p < 0.001). 10 to 19 replicates were carried out *per* treatment.

The qualitative and quantitative analysis of the electromyograms (EMGs) revealed different feeding patterns according to the treatment and feeding solution offered to each group **(Fig. 4B, C).** Regular and highly frequent pumping pulses, where each pump pulse represents a stroke of fluid being pumped by the muscles, can be observed for the uninjected and unrelated control (dsRNA-ctrl) groups, and the dsRNA-Rproppk28 group. Moreover, insects of these 3 groups were fully engorged (**Fig. S2**). The dsRNA-Rproppk014276 group presented fewer pumping pulses instead (**Fig. 4B, C**) (K-W = 19.6, p = 0.0006, Dunn’s *post hoc* comparisons against the uninjected group, p < 0.05), and insects did not engorge completely at the end of the experiment (**Fig. S2**). Interestingly, a similar and non-significant feeding performance was observed between uninjected insects offered with a solution lacking NaCl (no-salt solution) and the dsRNA-Rproppk014276 group offered with the optimal-salt solution (**Fig. 4B, C, S2**). These results indicate that salt detection at the PO is crucial to elicit feeding, since feeding is prevented if an insect does not sense salt. This response can be due to its absence or to the disruption of a salt-sensitive gene (*Rproppk014276*). We found that suppression of *Rproppk014276* expression, but not of *Rproppk28*, prevented NaCl detection and consequently feeding is hampered.

## Discussion

Pharyngeal taste machinery represents the last instance of food quality evaluation during ingestion. It was demonstrated that external and internal taste inputs are not functionally redundant; instead, both are required to control feeding in both non-blood feeders like *D. melanogaster* (Chen et al., 2019) and blood feeders like *R. prolixus* (Pontes et al., 2014). In this work, we characterized the PO, the internal taste organ of *R. prolixus*, through a multi-approach strategy. Thus, we examined different aspects of its neuroanatomy, physiology and genetics to unveil the role of this gustatory organ. PO sensilla house GRNs that reach the brain through the labral nerves. Gustatory information from PO-GRNs arrives at 2 regions of the SEZ where it will be primarily processed, although some inputs continue beyond this neuropile. Furthermore, the profile of gene expression in the PO provided molecular candidates likely involved in the gustatory assessment of the incoming blood. The knock-down of *Rproppk014276* gene expression showed its crucial role in feeding behavior, probably by affecting NaCl detection, one of the main phagostimulants of this species.

### The PO-GRNs are responsible for internal gustatory detection

The histological inspection of the short-peg sensilla on the PO of *R. prolixus* revealed the existence of sensory neurons. The morphology of internal gustatory organs was mostly studied in dipterans. For example, the PO of the fruit fly *D. melanogaster* and their taste sensilla have been deeply examined recently (Chen and Dahanukar, 2017). The cibarial organ of the mosquito *Ae. aegypti* and the tsetse fly *Glossina austeni*, equivalent to the PO in *R. prolixus*, hold taste sensilla which have also been postulated to be sensors of blood components (McIver and Siemicki, 1981; Rice et al., 1973). Although denominated differently, some of the morphotypes of sensilla present in the cibarial organ of both blood feeders are also innervated by chemosensory neurons (McIver and Siemicki, 1981; Rice et al., 1973).

We also reported the first physiological examination of the PO sensory neurons in a blood-feeding insect. Here, by registering from the PO, we demonstrated the neural response to relevant gustatory stimuli for *R. prolixus*. We obtained multicellular responses to two phagostimulants or appetitive compounds, NaCl and ATP, and to one aversive compound, caffeine. NaCl, a main constituent of blood, is necessary to all blood-feeding insects studied so far (e.g. *R. prolixus, Triatoma infestans, Ae. aegypti*, *Anopheles spp*, *Culex pipiens*, *Culiseta inornata*, *Glossina spp*., *Tabanus nigrovittatus*, *Simulium venustrum*, *Stomoxys calcitrans*, *Xenopsylla cheopis*) to achieve a normal and complete feeding response (Barrozo et al., 2022). Adenosine nucleotides, such as ATP and ADP, released from the erythrocytes and platelets upon shear stress on blood capillaries, also elicit gorging in most blood feeders (Barrozo, 2019). In contrast, detection of alkaloids in the ingested food, such as caffeine, quinine, theophylline, and berberine, drives the ultimate decision to reject a meal in *R. prolixus* (Pontes et al., 2014; Asparch et al., 2016; Muñoz et al., 2020). Similarly, quinine and caffeine also cause feeding avoidance of sucrose solutions in *Ae. aegypti* (Ignell et al., 2010), while denatonium benzoate and berberine negatively affect sugar feeding in *An. gambiae* (Kessler et al., 2013). But still, taste detection of aversive molecules in blood feeders is by far insufficient with respect to phytophagous insects (Barrozo, 2019). We showed that gustatory evaluation of appetitive and aversive molecules begins in GRNs located in the PO. The fine discrimination of the gustatory information will then occur in the brain, which will ultimately send an output message to the appropriate muscles to avoid or ingest a given resource.

### The subesophageal ganglion receives inputs from the PO

Previous morphological studies of pharyngeal neuron projections showed disparate results. In *D. melanogaster*, neurons in the pharynx project exclusively to the SEZ (Kwon et al., 2014). In turn, *Ae aegypti* sensory neurons of the cibarium project to the SEZ but also to the tritocerebrum whereas *G. austeni* neurons project exclusively to the tritocerebrum (Rice et al., 1973; McIver and Siemicki, 1981; Ignell and Hansson, 2005). Our results show that the axons of the PO-GRNs project through the labral nerves primarily into the SEZ, whereas some fibers continue to posterior ganglions. Furthermore, two different types of neurons reached two different regions of the SEZ, showing a differential topography of the PO-GRNs in the brain. This could be the basis of a differential processing of the information that comes from the pharyngeal organ, for example a sensory segregation of aversive inputs and appetitive as occurs in *Drosophila* (Harris et al 2015) and in mammals (Carleton et al., 2010). However, this hypothesis needs to be confirmed by functional studies.

### The PO transcriptome: candidate genes involved in feeding decisions

Our results showed a relevant compilation of sensory gene families in the PO tissue with the novel finding of *opsins* and *takeouts* (TOs) genes in the gustatory tissue of a blood-sucking insect. This is the first attempt to recognize molecular sensors in an exclusive gustatory tissue of a blood feeder and in particular in a triatomine insect. The current state of knowledge shows that *R. prolixus* engorges when ATP and low salt are present in the feeding solution, and rejects it when bitters, high or no salt are present (Barrozo, 2019). The underlying mechanisms of these opposing behaviors are still unknown and the first step towards their elucidation is to identify the molecular candidates present in the PO.

High expression of *RproGr26, RproGr27* and *RproGr28* was observed in the PO, as was also found in *R. prolixus* antennae previously (Latorre-Estivalis et al., 2017, 2022). The antennal expression pattern of these GRs was high and conserved in nymphs and adults suggesting a putative role in behaviors associated either to host search or interactions with congeners. Although finding these highly expressed genes in the PO allows us to contemplate this small set of GRs as major receptors in feeding events. Aversive feeding responses were described with exogenous chemicals for kissing bugs such as bitter compounds (Pontes et al., 2014). Molecular characterization of the pharyngeal organ of insects is scarce, however, this was performed for *D. melanogaster* in a more recent study which the expression of bitter and sugar GRs in the pharyngeal GRNs was reported, among other genes (Chen and Dakanukar, 2017). Among the repertoire of expressed GRs of *R. prolixus*, only *RproGr1* shows homology to *DmelGr43a*, which functions as a fructose receptor that triggers satiety responses once high fructose levels are detected (Miyamoto et al., 2012; Latorre-Estivalis and Lorenzo, 2019). Interestingly, *DmelGr43a*, which is expressed in a few neurons in the protocerebrum, appears to be restricted to some pharyngeal and tarsal GRNs in the fly gustatory system (LeDue et al., 2014). *R*. *prolixus* is an obligate blood feeder and fructose is not a common component found in blood, however, a recent report of plant feeding in these insects suggests that phytophagy could be an additional feeding habit (Díaz-Albiter et al., 2016). Our results reinforce these studies since a fructose sensor could be functional if these hematophagous insects would practice facultative phytophagy as is seen in mosquitoes (Barredo and DeGennaro, 2020; Jové et al., 2020). Fructose detection, as in the case of bitter detection, could represent the conservation of traits of past ancestors that were either predators or phytophagous (Pontes et al., 2014; Lehane, 2005). In either case, the feeding response to sugars in kissing bugs remains to be uncovered.

Alternatively, some PO-GRNs may evaluate characteristics other than palatability, such as temperature, humidity and even viscosity. The most expressed IR in the PO was the *RproIr93a*. This IR is a highly conserved among arthropods (Croset et al., 2010). In *D. melanogaster*, it was demonstrated to have a role in temperature preference in larvae and moisture detection in the adult antennae (Knecht et al., 2016). Since evidence shows that this receptor can participate in different mechanisms depending on which IRs are partnered with, functional hypotheses in the pharynx of *R. prolixus* are still indefinite. *RproIr75a* also presented enhanced expressions in the PO. In *D. melanogaster*, antennal *Ir75a* are tuned to acidic molecules, more specifically to acetic acid (Pietro-Godino et al., 2016). Because *R. prolixus* avoids feeding on acidic solutions (Barrozo pers. observation), *RproIr75a* in the PO would likely play a role in acidic molecule detection.

PPKs have been described to allow the detection of environmental stimuli like water, salts, or odors (Latorre-Estivalis et al., 2021). Enhanced expression was observed for the PPK*s Rprc000048, Rprc014276 (Rproppk014276*) and *Rproppk23* in the PO. Evidence for orthologues of *Rprc000048* is unreported. Recently, it was demonstrated that knocking-down the expression of *Rproppk28* and *Rproppk014276*, through RNA interference, resulted in a loss in avoidance of high concentration of salt (Pontes et al., 2022). It is plausible then, that *Rproppk014276* works as a salt sensor during gustatory assessment in the PO. Our RNAi experiments showed that the disruption of *Rproppk014276* expression prevented ingestion of an appetitive salt solution (see below).

TRPs are widely known for multimodal detection of stimuli and also for being highly conserved across different taxons (Matsuura et al., 2009; Peng et al., 2015). The PO repertoire of TRPs, although small, represents candidates for vital roles related to heat, water and noxious compounds detection as was previously demonstrated for other insect models (Gong et al., 2004; Bautista et al., 2005; Liu et al., 2007; Kim et al., 2010; Kang et al., 2010; Corfas and Vosshall, 2015). High levels of expression of *RproPainless* and *RproWaterwitch* are observed in the PO as was previously obtained for the antennal tissue (Latorre-Estivalis et al., 2017; 2022). The current state of knowledge shows that *Painless* and *Trpa1* are necessary to trigger aversive feeding responses to reactive electrophiles, such as allyl isothiocyanate (AITC), a pungent component of horseradish (Al-Anzi et al., 2006; Kang et al., 2010; Mandel et al., 2018). Moreover, *DmelPainless* intervenes in the avoidance of noxious heat, mechanical stimulation and dry environments (Tracey et al., 2003; Neely et al., 2011; Hwang et al., 2012). *D. melanogaster* flies detect moist air with their antennae using Waterwitch (Liu et al., 2007). Likely, *RproPainless* is an appropriate candidate to sense noxious heat and chemicals, whilst *RproWaterwitch* could fulfill a role in water sensing during feeding in the PO.

The novel finding of opsins in the PO of *R. prolixus* is remarkable since recently it was demonstrated their role in *D. melanogaster* taste sense (Leung et al., 2020). Moreover, it was demonstrated that opsins sense bitter tastants through a signal cascade that also includes a Gq protein, a *phospholipase C* and a *Trpa1*. This shows that not only a novel protein is involved in the insect taste sense but also a reminiscent mechanism of mammalian taste transduction such as a GPCR signaling pathway. *R. prolixus* opsins, but especially the *RproRh7*, could be interesting candidates to unveil their role in the detection of bitter compounds in these insects.

High expression of several OBPs, CSPs and TOs was also found in the PO. Our data support the current knowledge that showed that insect OBPs and CSPs are highly expressed in taste sensilla (Leal, 2013; Sánchez-Gracia et al., 2009). Enhanced expression of these auxiliary proteins could indicate a major role during blood feeding. They have been reported to act as solubilizers and carriers of hydrophobic nutrients in the mouthparts of blowflies, moths and butterflies, and as surfactants to reduce pressure during sucking and as detectors of nutrients in moths (Nagnan-Le Meillour et al., 2000; Ishida et al., 2013; Liu et al., 2014; Zhu et al., 2016). Starvation seems to regulate the expression of TO genes in *D. melanogaster* and food intake was shown to be indirectly affected by this gene family, as was demonstrated for mutant flies (Meunier et al., 2007). Our results showed that the TO family is the most expressed in the PO, suggesting putative feeding regulatory functions in *R. prolixus*.

The high expression of serotonin, acetylcholine and dopamine receptors suggests relevant functions of these neurotransmitters in the PO functioning. In several insect models, a strong link between biogenic amine levels with modulation during hunger/satiety has been demonstrated. Serotonin was demonstrated to modulate appetite, usually depressing feeding in flies, bees, ants and other insect models (Blenau and Erber, 1998; Dacks et al., 2003; Haselton et al., 2009; Falibene et al., 2012; Schoofs et al., 2018). In *R. prolixus*, injection of the neurotoxin 5,7-DHT, which depleted serotonin of peripheral neurons, led to a decrease in blood intake (Cook and Orchard, 1990). Dopamine activates *DopEcR* in the GRNs to enhance sensitivity to sugar in hungry flies and 20-hydroxyecdysone activates this receptor to repress feeding and promote pupation in lepidopteran insects (Inagaki et al., 2012; Kang et al., 2019). The presence of acetylcholine receptors in the alimentary canal is novel and its putative role is unknown. The predominant presence of neurotransmitter receptors in the PO reinforces this organ as a site for the modulation of food acceptance in relation to the internal state and needs of the insect.

The neuroendocrine system regulates feeding behavior and food search in gustatory neural circuits (Nässel and Zandawala, 2019). The *RproGPA2/GPB5* heterodimer was highly expressed, for both the neuropeptide precursor and its receptor, in the PO. The GPA2/GPB5 system was suggested to participate in the control of ion and water balance in the hindgut of *Ae. aegypti* (Paluzzi et al., 2014). *RproAst-CC*was the highest expressed NP precursor gene in the PO, as previously found in *R. prolixus* antennae, pointing to a relevant role in the regulation of the perception of chemical stimuli (Latorre-Estivalis et al., 2020). Other highly expressed precursor genes in the PO were the *RproITG-like, RproITP, RproMIP, RproIDLSRF-like* peptide and *RproPDF*. The abundance of *RproITG-like* peptides is modulated 24 h post-blood meal (Sterkel et al., 2011). *MIP* reduces the sensitivity toward food in *D. melanogaster;* its silencing causes an increase in food intake and body weight, inducing satiated flies to behave like starved ones (Min et al., 2016). The NPRs with the highest expression in the PO were the *RproFalp/ProcR, RproCT/DhR1, RproCCAPR1, RproCRF/DHR2, RproNPLPR* and *RproInR*. Proctolin stimulates contractions in the midgut and hindgut of insects, suggesting a similar function in the PO of *R. prolixus* (Orchard et al., 2011). *RproCT/DhR1* is also related to feeding modulation since it increases the contractions of salivary glands and hindgut (Brugge et al., 2008; Zandawala et al., 2013). Besides, the regulation of blood feeding by the *RproCCAP* system is reinforced by the expression of *RproCCAP* in salivary glands and the *RproCCAP* precursor in projections arriving at the salivary glands (Lee et al., 2013). Insects injected with synthetic *RproCRF/DH* peptide before feeding ingested a significantly reduced blood meal (Mollayeva et al., 2018). Moreover, *RproNPLP1* is modulated in response to a blood meal in *R. prolixus* (Sterkel et al., 2011). The expression profiles of NPs and NPRs detected in the PO reinforces the hypothesis of these poorly studied neuropeptide systems in regulating *R. prolixus* feeding.

### *Rproppk014276* is responsible for salt sensing in the PO

Since salt concentration is an important factor modulating feeding decisions, the implications of PPKs on salt sensing make them interesting candidates for functional genetic studies. Highly concentrated salt and no-salt solutions give rise to aversive behaviors whilst optimal (appetitive) concentration of salts in combination with ATP elicit feeding (Pontes et al., 2017). Knocking down the expression of two PPKs, previously related to salt detection (Pontes et al., 2022), allowed us to establish the functionality of the PO during food assessment. We demonstrated that RNAi knockdown of *Rproppk014276* interferes with feeding acceptance. The role of *Rproppk28* in salt sensing could not be revealed under the experimental conditions of this work. Certainly, its function in the PO needs further investigation. Beyond the different functions that the large repertoire of genes expressed in the PO can have, our main objective was to establish the crucial role of the PO in food evaluation. Similarly, pharyngeal neurons control food choice and intake in *D. melanogaster*, as was demonstrated in *Poxn* mutant flies (Chen et al., 2019). In the case of *R. prolixus*, the PO is the main gustatory organ in the pharynx involved in food evaluation. Interestingly, *Rproppk014276* modulates the aversive response to high concentration of salt in the antennae, whereas it modulates the feeding acceptance to optimal (appetitive) salt level in the PO (Pontes et al., 2022). Likely, the differential chemosensory context of the antennae and the PO determines the salience of this gene’s activation.

## Conclusions

The PO, located in the first portion of the alimentary canal, constitutes the last instance of food evaluation in *R. prolixus*. Due to the difficulty associated with size and physical access, the contribution in feeding by internal taste organs has been little studied despite their great relevance across the animal taxa. This work shows for the first time that the PO of *R. prolixus* presents neurons, within the short-peg sensilla, connected to higher brain centers and expression of genes with the potential function of detecting the different chemical and physical properties of blood. Our characterization of the PO is the first step towards elucidating blood acceptance and rejection mechanisms in a non-traditional blood-sucking insect model, while also envisioning extrapolation to other blood-sucking models, such as mosquitoes.

## Methods

### Animals

Fifth-instar nymphs and adults of *R. prolixus* were used throughout the experiments. The rearing conditions of insects were 28 °C, ambient relative humidity and 12h: 12 h light: dark cycle. Insects were kept unfed following ecdysis. Fifth-instar larvae 7 - 21 days post-ecdysis were used for electrophysiology, feeding assays and RNA-Seq. Adults used for neuroanatomy experiments were 7 - 9 day-old post-ecdysis. Feeding experiments were carried out at the beginning of the insects’ scotophase, the moment of the day that the kissing bugs had a maximal motivation to feed (Barrozo et al., 2017).

### Neuroanatomy

#### Histology

For the histological analysis of the PO, the heads of insects were fixed for 3 h in a mixture of 2.5% glutaraldehyde and 2.0% paraformaldehyde in phosphate buffer (pH 7.3) with glucose and CaCl_2_. After dehydration, they were embedded *via* propylene oxide in Durcupan ACM (Electron Microscopy Sciences, Pennsylvania, US). Blocks were serially sectioned at 5 - 10 μm using glass knives mounted in a microtome. The sections were stained on a hot plate with Methylene Blue and observed under a light microscope (Olympus, JP).

#### Anterograde fills

Live insects were dorsally fixed to a glass support. An opening was made in the cuticle of the head in the antero-dorsal region of the rostrum, in order to reach the somas and axons of the PO-GRNs. Subsequently, a drop of distilled water was applied for 6 min. Following this time, the distilled water was absorbed and a drop of the neuronal tracer rhodamine (1% in distilled water, Dextran, Tetramethylrhodamine, 3000MW, Anionic, Lysine Fixable, Thermo Fisher, Massachusetts, US) was applied and covered with vaseline to avoid dehydration in a total of 28 adult insects. Then, live insects were maintained inside closed Petri dishes with wet cotton (to assure the maintenance of a humid ambient) for 48 h to allow the neuronal tracer to diffuse to the brain at 8°C. After this time, the brains were dissected in Millonig’s buffer and fixed in 4% paraformaldehyde overnight at 4°C. Then, brains were rinsed in Millonig’s buffer, dehydrated through sequential ethanol series and finally cleared and mounted in methyl salicylate (Pontes et al., 2022). Whole mounts were optically sectioned and scanned with a laser scanning confocal microscope (Olympus FV300/BX61, Centro de Microscopía Avanzada, Facultad de Ciencias Exactas y Naturales, Universidad de Buenos Aires, Buenos Aires, AR).

### Electrophysiological recordings

Global action potentials of the PO-GRNs were recorded through extracellular recordings. To get access to the PO, live insects were fixed dorsally and a first opening was made in the cuticle of the head in the anterior-ventral region of the rostrum. A second opening of the ventral wall of the pharynx was made to finally expose the PO. An Ag/AgCl reference electrode was located inside the insect’s abdomen, while a tungsten recording electrode was placed next to PO. One hundred μl of the following gustatory stimuli were applied to the preparation: 0.15 M NaCl, 1 mM ATP (in 0.15 M NaCl) and 5 mM caffeine (in 0.15 M NaCl). Each insect was first stimulated with NaCl, and 1 min later with either ATP or caffeine. The preparation was rinsed with 0.15 M NaCl between stimulations. The interval between subsequent stimulations was 5 min. The biological signals were amplified, filtered (gain ×10, TastePROBE DTP-02, Syntech; gain × 100, eighth-order Bessel, pass-band filter: 10 - 3000 Hz, Dagan Ex1), digitized (Data Translation DT9803; sampling rate: 10 kHz, 16 bits) and stored in a PC. Spike detection and firing frequency quantification were performed off-line using dbWave software (Marion-Poll, 1996). ATP was purchased at Sigma-Aldrich (St Louis, US) and caffeine at Biopack (Buenos Aires, AR).

The responses of PO-GRNs to the 3 gustatory stimuli were statistically evaluated using the Kruskal-Wallis test (α = 0.05). Subsequently, the responses to ATP and caffeine were compared to the response to NaCl through Dunn’s *post hoc* comparisons. 9 to 14 replicates were carried out.

### RNA extraction

A total of 150 POs were dissected, removing them with forceps and immediately placing them in 3 separate tubes (50 POs each) added with RNAlater™ stabilization solution (Invitrogen, Carlsbad, USA) and kept at −80 °C. Tissue samples were later transferred onto a Whatman™ filter paper Cat N° 1001–055 (Whatman Inc., Florham Park, US) using forceps at room temperature. After absorption of the solution excess, the tissues were immersed in 200 μL of TRIzol^®^ reagent (Invitrogen, Carlsbad, US), macerated with a pestle, and 800 μL of TRIzol was added. The RNA extraction was performed as described in Bellantuono et al., 2012. The total RNA was eluted from the columns with 50 μL of RNase-free water, and an aliquot from each sample was separated for RNA quality check using Agilent 2100 Bioanalyzer (Agilent Technologies, Palo Alto, US). The RNA samples were immediately stored at −80 °C for further analysis.

### Library preparation and RNA-seq

RNA library preparations and sequencing reactions were conducted at GENEWIZ, LLC. (South Plainfield, US), accordingly to the following protocol. RNA samples were quantified using Qubit 2.0 Fluorometer (Life Technologies, Carlsbad, US) and RNA integrity was checked using Agilent TapeStation 4200 (Agilent Technologies, Palo Alto, US). RNA sequencing libraries were prepared using the NEBNext Ultra RNA Library Prep Kit for Illumina using the manufacturer’s instructions (NEB, Ipswich, US). Briefly, mRNAs were initially enriched with Oligod (T) beads. Enriched mRNAs were fragmented for 15 minutes at 94 °C. First-strand and second-strand cDNAs were subsequently synthesized. cDNA fragments were end-repaired and adenylated at 3’ ends, and universal adapters were ligated to cDNA fragments, followed by index addition and library enrichment by PCR with limited cycles. The sequencing library was validated on the Agilent TapeStation, and quantified by using Qubit 2.0 Fluorometer as well as by quantitative PCR (KAPA Biosystems, Wilmington, US). The sequencing libraries were clustered on a flow cell. After clustering, the flowcell was loaded on the Illumina HiSeq instrument (4000 or equivalent) according to the manufacturer’s instructions. The samples were sequenced using a 2×150bp paired-end configuration. Image analysis and base calling were conducted by the HiSeq Control Software (HCS). Raw sequence data (.bcl files) was converted into fastq files and de-multiplexed using Illumina’s bcl2fastq 2.17 software. One mismatch was allowed for index sequence identification. The raw sequence dataset is available in the NCBI BioProject database with the accession number BioProject ID PRJNA674000.

### Bioinformatic analysis

The presence of Illumina sequencing adapters and the quality of reads from the sequencing facility were analyzed using the FASTQC tool (www.bioinformatics.babraham.ac.uk/projects/fastqc). Then, Illumina adapters and those bases from 5’ and 3’ ends with quality scores lower than 5 (TRAILING: 5 and LEADING: 5 parameters) were eliminated from the reads using Trimmomatic v0.32 in the paired-end mode (Bolger et al., 2014). The SLIDING-WINDOW parameter was fixed at 4:15 and only those reads longer than 50 bp were kept for the next steps. Afterward, trimmed and cleaned reads were mapped and counted to the *R. prolixus* genome assembly (version RproC3.3 from VectorBase) through STAR v.2.6.0 (Dobin et al., 2013) with default parameters and an edited General Feature Format (GFF) file generated by Latorre-Estivalis et al. (2020). Raw counts are included in the **Supplementary Table 1**. Heatmaps showing gene expression (expressed as Log_10_(TPM+1)) of the different protein families were prepared using the gplot package in R (R Core Team 2020) (**Supplementary Table 2**). TPM values less than 1 were considered as genes not abundantly expressed in the PO or below detection limits.

### RNA interference

Double-strand RNA of *Rproppk28* and *Rproppk014276* were synthesized and injected as described in Pontes et al. (2022). PCRs were carried out by using specific primers conjugated with 20 bases of the T7 RNA polymerase promoter. The beta-lactamase gene (β-lact) of *Escherichia coli*, an ampicillin resistance gene, was also amplified from the pBluescript SK plasmid as control of dsRNA injection. PCR products for *Rproppk014276, Rproppk28* and β-lact were used as templates for dsRNA synthesis using the MEGAscript™ RNAi Kit (Thermo Fisher Scientific, Massachusetts, US). All PCR fragments were sequenced and checked for their similarity to the expected fragments. After synthesis, the purity and integrity of dsRNA were confirmed through a 1.5 % agarose gel and quantified using NanoDrop™ (Thermo Fisher Scientific, Massachusetts, US). The specificity of the dsRNA for each PPK was validated *in silico* using BLASTn searches against the *R. prolixus* genome sequences, and each dsRNA sequence showed a unique and complete hit against its target sequence.

Insects were randomly separated into 4 experimental groups. Two groups of insects were injected with the corresponding dsRNAs: dsRNA-Rproppk014276 and dsRNA-Rproppk28. A third group was injected with the dsRNA of β-lact, representing a control for dsRNA injection (dsRNA-ctrl). A microliter syringe (World Precision Instruments, Florida, US) was used to inject 2 μl of dsRNA (concentration = 1.25 μg/μl) diluted in PBS 1X into the thorax. The fourth group of insects was maintained intact (uninjected group) as an additional control. Eleven days after dsRNA injection, insects of each group were tested for RT-qPCR verification of gene expression knock-down or behavioral assays (see below).

Knock-down efficacy was checked and described in Pontes et al. (2022). Statistical differences between the transcript levels of the experimental groups were analyzed using the Kruskal-Wallis test and *post hoc* Dunn’s multiple comparison test (α = 0.05) **(Fig. S1)**.

### Feeding response of dsRNA injected and control insects

The activity of the pharyngeal and cibarial muscles involved in feeding was recorded in dsRNA injected and uninjected insects for 10 minutes. For this, an artificial feeder coupled to an electromyographic recording system was used (Pontes et al., 2017). Briefly, the device consists of two containers, the feeder and the insect container. The feeder contains the feeding solution, and the insect container, which is attached to the feeder, holds the insect. The feeder is covered with a latex membrane that mimics the host skin, which insects can easily pierce to access the feeding solution. A solution of 1 mM ATP in 0.15 M NaCl was used as the feeding solution for all groups, which constitutes an appetitive optimal-salt solution for *R. prolixus* (Pontes et al., 2017). In addition, a solution of 1 mM ATP in distilled water was also tested in a control group of uninjected insects, which constitutes a non-appetitive salt-free solution (Pontes et al., 2017). A metallic mesh connected to a copper wire (recording electrode) was placed inside the insect container. The insect used this metallic mesh to climb and access the feeder. A silver wire (reference electrode) was located inside the feeder in contact with the feeding solution. Both electrodes were connected to a differential amplifier (HotBit HB3600, DE). The experiment started once the insect inserted its mouthparts into the feeder, thus, the electrical circuit was closed generating a base conductance. The changes in the baseline signal were attributed to contractions produced by the sucking muscles when the insect began to feed. Recorded signals were amplified (gain x200) and digitized with aid of the A/D converter of the oscilloscope (Tektronix TDS 210, Oregon, US) connected to a PC. Recordings were acquired and analyzed off-line using software designed *ad hoc* (Diego Anfossi, Héctor Salas Morales, *unpubl.*). The pumping frequency of muscles was calculated by counting the number of pumps (i.e. each pump or peak represented a muscle contraction) during the time the insect sucked the solution. The weight gain of insects during feeding was also registered, and a feeding factor was calculated as (Wf - Wi)/Wi; where Wf: final weight, Wi: initial weight **(Fig. S2)**. Statistical differences among experimental groups were assayed using the Kruskal-Wallis test (α = 0.05). *A posteriori* comparisons were carried out between the uninjected group (fed on optimal-salt solution) and the rest of the groups using Dunn’s *post hoc* comparisons. 10 to 19 replicates were carried out.

## List of abbreviations

PO: pharyngeal organ;
PO-GRNs: pharyngeal organ gustatory receptor neurons;
SEZ: subesophageal zone;
ds-RNA: double-stranded RNA;
TPM: Transcripts *Per* kilobase *per* Million read;
GRs: gustatory receptors;
IRs: ionotropic receptors;
PPKs: pickpocket channels;
TRPs: transient receptor potential channels;
Ops: opsins;
OBPs: odorant-binding proteins;
CSPs: chemosensory proteins;
TOs: takeouts;
NPs: neuropeptide precursors;
NPRs: neuropeptide precursor receptors;
BARs: biogenic amines receptors.

## Declarations

### Ethics approval and consent to participate

Not applicable

### Consent for publication

Not applicable

### Availability of data and materials

All data needed to evaluate the conclusions in the paper are present in the paper and/or the supplemental information. All data will be deposited on Mendeley Data (https://data.mendeley.com/datasets/) upon acceptance of the manuscript.

Any additional information is available from the lead contact upon request.

### Authors’ contributions

RBB designed the research. IOI, RBB, JLE, AC, MBA, HSM, GP, ACS, TI performed the experiments. IOI, RBB, JLE, SO, ACS, MBA, MDG analyzed the data. IOI, RBB, JLE, SO, ACS, MDG wrote the paper. All authors revised the manuscript. All authors read and approved the final manuscript.

## Acknowledgements

We thank Marcelo Lorenzo for the discussions. We also thank Claudio Lazzari for providing the facilities to process the histological tissue samples. We thank the CONICET, UBA and ANPCyT.

## Supplementary Figures and Tables

**Figure S1.** Expression levels of Rproppk014276 and Rproppk28 after dsRNA injection. Bars represent the relative expression levels (mean ± s.e.m.) of **(A)** Rproppk014276 and **(B)** Rproppk28 in the four experimental groups. The transcript levels of Rproppk014276 and Rproppk28 in the antennae were significantly decreased to the corresponding control groups. Selective knock-down of the gene of interest was also checked for both genes (last bars). Asterisks indicate significant differences across groups (Kruskal-Wallis test, Dunn’s *post hoc* comparisons, p < 0.001). 6 replicates were carried out *per* treatment. Data from Pontes et al. (2022).

**Figure S2**. Weight gain of treated insects during feeding on the optimal salt or on the salt-free (no salt) solutions. No differences were found between uninjected, dsRNA-ctrl and dsRNA-Rproppk28 groups. dsRNA-Rproppk014276 insects, however, showed significantly lower weight gain than the other groups. The dsRNA-Rproppk014276 group offered with optimal salt showed no differences from uninjected insects fed on a salt-free solution. Scatter plots are shown and bars represent the mean feeding factor (mean ± s.e.m). The feeding factor was calculated as a normalized weight gain as follows (Wf - Wi)/Wi; where Wf: final weight, Wi: initial weight. Asterisks indicate significant differences of each group to uninjected insects fed on the optimal-salt solution (Kruskal-Wallis test, Dunn’s *post hoc* comparisons, p < 0.0001). 10 to 20 replicates were carried out *per* treatment.

**Supplementary Table S1**. Raw count matrix generated by STAR after mapping trimmed and cleaned reads against the *R. prolixus* genome.

**Supplementary Table S2.** Transcripts *per* kilobase *per* Million read (TPM) values in the three replicates for all targeted genes. TPM values > 1 are in light blue.

