## Supplementary figures and images for "Food recognition in a blood-feeding insect: characterization of the pharyngeal taste organ"

### Figure S1

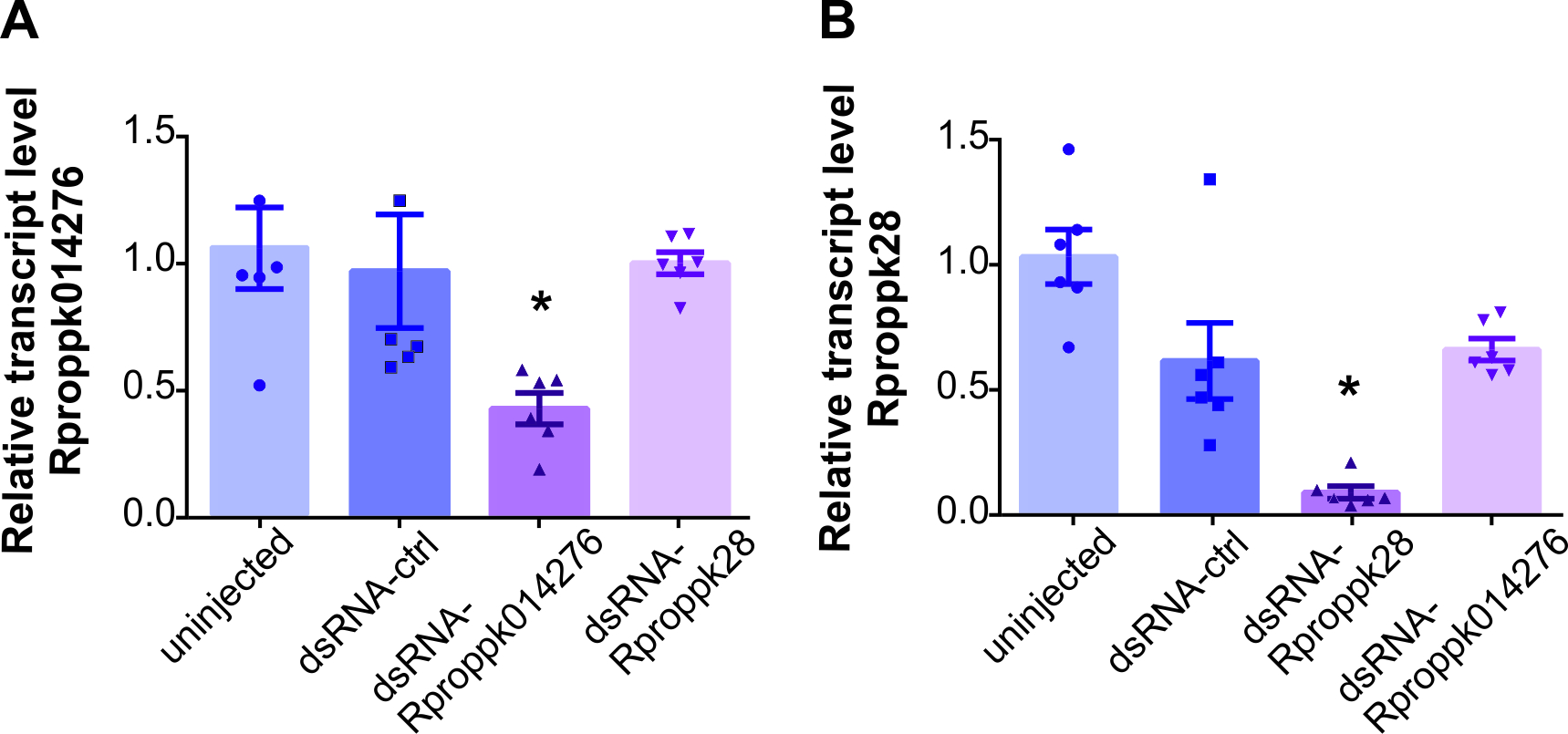

### Figure S2

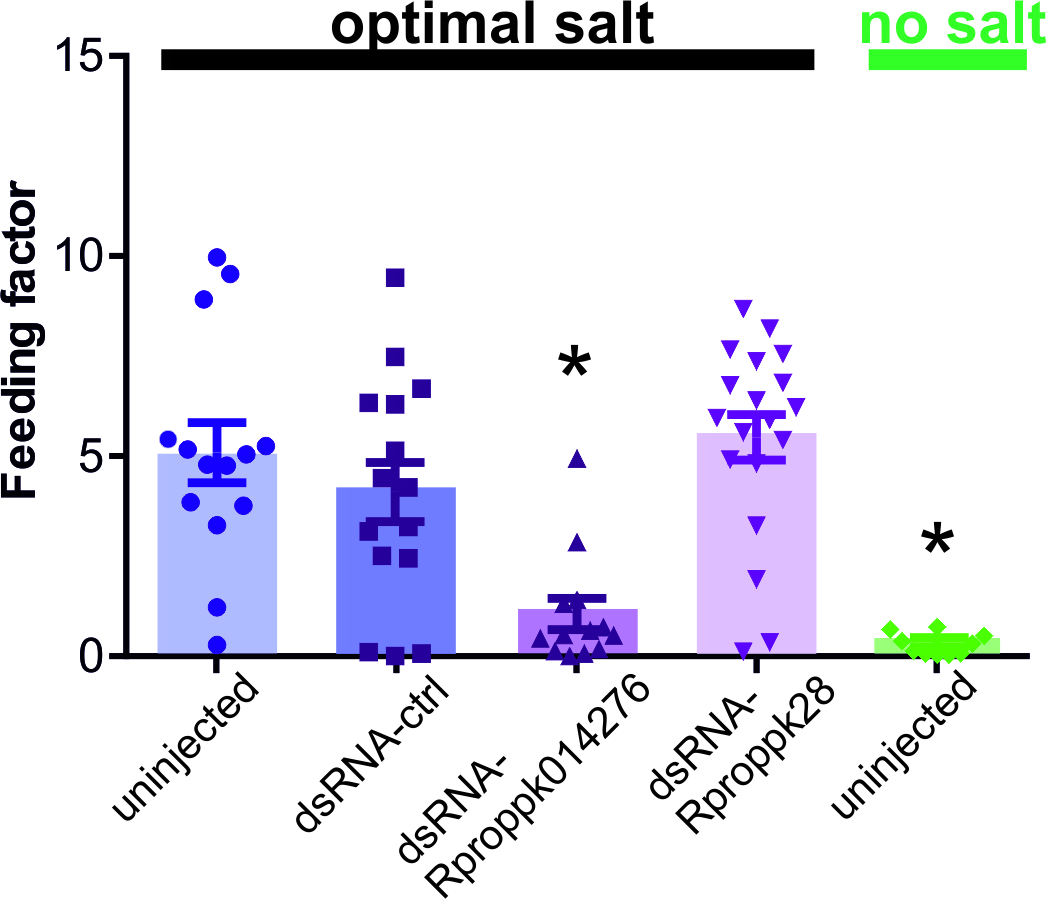
